# Dissecting the genetics of complex traits using summary association statistics

**DOI:** 10.1101/072934

**Authors:** Bogdan Pasaniuc, Alkes L. Price

## Abstract

During the past decade, genome-wide association studies (GWAS) have successfully identified tens of thousands of genetic variants associated with complex traits and diseases. These studies have produced vast repositories of genetic variation and trait measurements across millions of individuals, providing tremendous opportunities for further analyses. However, privacy concerns and other logistical considerations often limit access to individual-level genetic data, motivating the development of methods that analyze summary association statistics. Here we review recent progress on statistical methods that leverage summary association data to gain insights into the genetic basis of complex traits and diseases.

## Introduction

Genome-wide association studies (GWAS) have been broadly successful in identifying genetic variants associated to complex traits and diseases, explaining a significant fraction of narrow-sense heritability and occasionally pinpointing biological mechanisms^1^. These studies have produced vast databases of genetic variation (typically at the level of common single nucleotide polymorphisms (SNPs) included on genotyping arrays) in millions of individuals across hundreds of complex traits. Further analyses of this data can yield important insights into the genetics of complex traits, but privacy concerns and other logistical considerations often restrict access to *individual-level data*. On the other hand, *summary association statistics*, defined here as per-allele SNP effect sizes (log odds ratios for case-control traits) together with their standard errors, are often readily available and can be used to compute *z-scores* (per-allele effect sizes divided by their standard errors; see Figure 1); we note that in some applications, allele frequencies may also be required. A partial list of publicly available summary association statistics from large GWAS is provided in Table 1. Summary statistics also offer advantages in computational cost, which does not scale with the number of individuals in the study. These advantages have motivated the recent development of many new methods for analyzing summary association data, often in conjunction with linkage disequilibrium (LD) information from a population reference panel such as 1000 Genomes^2^.

**Figure 1.**
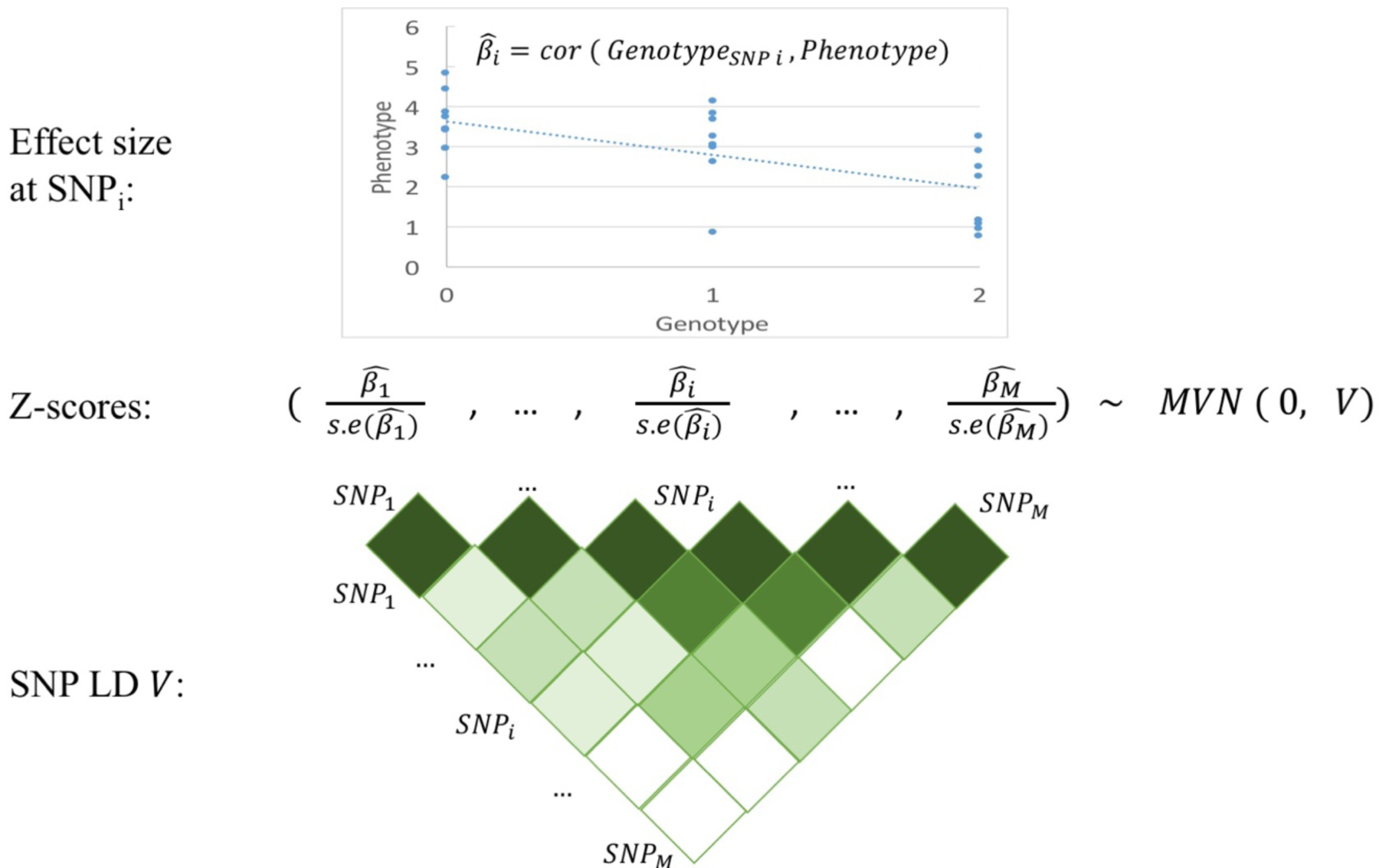
Illustration of summary association statistics. Per-allele SNP effect sizes (and their standard errors) are typically estimated by regressing the phenotype on the genotype values at the SNP of interest (top). At large sample sizes, the vector of z-scores (effect sizes divided by their standard errors) at a locus are approximated by a multivariate normal distribution with mean 0 and variance equal to the LD matrix *V* (bottom).

**Table 1.** Publicly available summary association statistics. We provide a partial list of publicly available summary statistics from GWAS with sample size at least 20,000. *: includes specialty chip data; not suitable for analysis using LD score regression and its extensions.

| **Trait** | **N** | **Reference** | **URL** |
| --- | --- | --- | --- |
| Age at menarche | 127,884 | Perry et al. 2014<br>Nature | http://www.reprogen.org/ |
| Alzheimer's | 54,162 | Lambert et al. 2013<br>Nat Genet | http://www.pasteur-lille.fr/en/recherche/u744/igap/igap_download.php |
| Bone mineral density | 53,236 | Zheng et al Nat Genet 2015 | http://www.gefos.org/?q=content/data-release-2015 |
| BMI | 122,033 | Speliotes et al. 2010<br>Nat Genet | http://www.broadinstitute.org/collaboration/giant/index.php/GIANT_consortium_data_files |
| BMI* | 322,154 | Locke et al. 2015<br>Nature | http://www.broadinstitute.org/collaboration/giant/index.php/GIANT_consortium_data_files |
| Coronary artery disease | 77,210 | Schunkert et al. 2011<br>Nat Genet | http://www.cardiogramplusc4d.org/ |
| Crohn's disease | 20,883 | Jostins et al. 2012<br>Nature | http://www.ibdgenetics.org/downloads.html |
| Crohn's disease* | 51,874 | Liu et al. 2015 Nat Genet | http://www.ibdgenetics.org/downloads.html |
| Depressive symptoms | 161,460 | Okbay et al. 2016<br>Nat Genet | http://ssgac.org/documents/ |
| Ever smoked | 74,035 | Furberg et al. 2010<br>Nat Genet | http://www.med.unc.edu/pgc/downloads/ |
| Fasting glucose | 58,074 | Manning et al. 2012<br>Nat Genet | http://www.magicinvestigators.org/downloads/ |
| HbA1C | 46,368 | Soranzo et al. 2010<br>Diabetes | http://www.magicinvestigators.org/downloads/ |
| HDL | 97,749 | Teslovich et al. 2010<br>Nature | http://www.broadinstitute.org/mpg/pubs/lipids2010/ |
| HDL* | 188,577 | Willer et al. 2013 NG | http://csg.sph.umich.edu/abecasis/public/lipids2013/ |
| Height | 131,547 | Lango Allen et al.<br>2010 Nature | http://www.broadinstitute.org/collaboration/giant/index.php/GIANT_consortium_data_files |
| Height* | 253,288 | Wood et al. 2014 Nat Genet | http://www.broadinstitute.org/collaboration/giant/index.php/GIANT_consortium_data_files |
| Hip Circumference | 213,038 | Shungin et al Nat Genet 2015 | http://portals.broadinstitute.org/collaboration/giant/index.php/GIANT_consortium_data_files |
| IBD (Crohn's/UC) | 34,652 | Jostins et al. 2012<br>Nature | http://www.ibdgenetics.org/downloads.html |
| IBD* (Crohn's/UC) | 65,643 | Liu et al. 2015 Nat Genet | http://www.ibdgenetics.org/downloads.html |
| LDL | 93,354 | Teslovich et al. 2010<br>Nature | http://www.broadinstitute.org/mpg/pubs/lipids2010/ |
| LDL* | 188,577 | Willer et al. 2013 NG | http://csg.sph.umich.edu/abecasis/public/lipids2013/ |
| Neuroticism | 170,911 | Okbay et al. 2016<br>Nat Genet | http://ssgac.org/documents/ |
| RA (Europeans) | 38,242 | Okada et al. 2014<br>Nature | http://plaza.umin.ac.jp/yokada/datasource/software.html |
| RA* (Europeans) | 58,284 | Okada et al. 2014<br>Nature | http://plaza.umin.ac.jp/yokada/datasource/software.html |
| RA (East Asians) | 22,515 | Okada et al. 2014<br>Nature | http://plaza.umin.ac.jp/yokada/datasource/software.html |
| Schizophrenia | 70,100 | Ripke et al. 2014<br>Nature | http://www.med.unc.edu/pgc/downloads/ |
| Subjective well-being | 298,420 | Okbay et al. 2016<br>Nat Genet | http://ssgac.org/documents/ |
| Triglycerides | 94,461 | Teslovich et al. 2010<br>Nature | http://www.broadinstitute.org/mpg/pubs/lipids2010/ |
| Triglycerides* | 188,577 | Willer et al. 2013 NG | http://csg.sph.umich.edu/abecasis/public/lipids2013/ |
| Type 2 diabetes | 60,786 | Morris et al. 2012<br>Nat Genet | http://www.diagram-consortium.org/ |
| Ulcerative colitis | 27,432 | Jostins et al. 2012<br>Nature | http://www.ibdgenetics.org/downloads.html |
| Ulcerative colitis* | 47,746 | Liu et al. 2015 Nat Genet | http://www.ibdgenetics.org/downloads.html |
| Waist Circumference | 232,101 | Shungin et al Nat Genet 2015 | http://portals.broadinstitute.org/collaboration/giant/index.php/GIANT_consortium_data_files |
| Waist Hip Ratio | 212,248 | Shungin et al Nat Genet 2015 | http://portals.broadinstitute.org/collaboration/giant/index.php/GIANT_consortium_data_files |
| Years of education | 328,917 | Okbay et al. 2016<br>Nature | http://ssgac.org/documents/ |

Here, we review these summary statistic-based methods. First, we review methods for performing single-variant association tests, including meta-analysis, conditional association and imputation using summary statistics. Second, we review methods for performing gene-based association tests by incorporating transcriptome reference data or aggregating signals across multiple rare variants. Third, we review methods for fine-mapping causal variants, including integration of functional annotation and/or trans-ethnic data. Fourth, we review methods for constructing polygenic predictions of disease risk and inferring polygenic architectures. Finally, we review methods for jointly analyzing multiple traits. We conclude with a discussion of research areas where further work on summary statistic based methods is needed.

### Single-variant association tests

#### Meta-analysis using fixed-effects or random-effects models

Large consortia often combine multiple GWAS studies into a single aggregate analysis to boost power for discovering SNP associations of small effect. Studies are combined either by jointly analyzing summary association results from each study (*meta-analysis*) or by re-analyzing individual-level data across all studies (*mega-analysis*)^3^. It has been shown that meta-analysis attains similar power for association as mega-analysis, with fewer privacy constraints and logistical challenges (since only summary association data is shared across studies)^4^. Meta-analysis is usually performed using fixed-effects approaches, which assume that true effect sizes are the same across studies. If true effect sizes are expected to differ across studies, this heterogeneity can be explicitly modeled using random-effects methods, which include an extra variance term in the model to account for heterogeneity. Traditional random-effects methods allow for heterogeneity under the null model, leading to low power even when heterogeneity is present. This motivated the development of a random-effects method based on a null model of no-heterogeneity, which increases power over traditional random-effects methods^5^. Under this framework, a statistical test against a null model of no-heterogeneity can be viewed as a summation of a fixed-effect component and a heterogeneity component, thus connecting fixed-effects and random-effects meta-analysis^5^. Subsequent work has introduced the concept of posterior probability for each study to have a non-zero effect, aiding interpretation and power when only a subset of studies have non-zero effect^6^.

#### Conditional association using LD reference data

Conditional association, in which the association between SNP and trait is evaluated after conditioning on the top SNP at a locus, can be used to identify multiple signals of association at a previously identified GWAS locus. Conditional association methods have traditionally required individual-level data in order to jointly fit multiple SNPs. Recent work has shown that conditional and joint association analysis of multiple SNPs can be approximated using only summary association statistics together with linkage disequilibrium (LD) information estimated from a population reference panel such as 1000 Genomes (see Box 1)^7^. This has enabled the discovery of new secondary associations at known loci for height, BMI, and other complex traits and diseases, increasing the variance explained by GWAS associations for these traits^8-10^; for example, in a recent height GWAS, approximate conditional analysis using summary data identified 697 genome-wide significant SNPs, including 34 SNPs with *r*^2^>0.1 to a more significant SNP at the same locus^8^.

#### Imputation using summary association statistics

A standard approach to boost association power in GWAS is to leverage LD information from a population reference panel to impute genotypes at variants not typed in the study^11^. Imputation is traditionally performed using individual-level data, which requires substantial computational resources and can be logistically cumbersome when new reference panels become available, particularly for large consortia combining data from multiple studies. As an alternative to imputation using individual-level data, approaches have been developed to perform imputation directly at the level of summary statistics^12-18^. The key insight of these approaches is that LD induces correlations between z-scores, which can be modeled using a multivariate normal (MVN) distribution with variance equal to the LD correlation matrix^19^. Thus, z-scores at untyped SNPs can be imputed from observations at typed SNPs using conditional means and variances of the MVN distribution. Imputation using summary statistics recovers >80% of the information from imputation using individual-level data at common variants^14-16^, and is practical and efficient since the imputed summary statistics are linear combinations of the observed statistics (see Box 1). However, imputation using summary statistics cannot capture non-linear relationships between SNPs, which are modeled using haplotypes in imputation from individual-level data.

Conditional association and imputation using summary statistics critically rely on accurate LD information from a population reference panel. Even in the best case where the reference population closely matches the GWAS population, the relatively small reference panel size (typically hundreds or at most thousands of individuals) makes accurate estimation of a large number of LD parameters a challenge. This motivates regularization of the estimated LD matrix, both to maximize accuracy and to ensure robustness in the case of imputation using summary statistics, as mis-estimation of the variance of imputed statistics can lead to false-positive associations. A simple approach to regularization is to set all correlations between distal SNPs to zero, based on a fixed distance threshold^7^ or approximately independent LD blocks inferred from the data^20^. An alternative is to specify a prior distribution and compute Bayesian posteriors^12^; data can be combined across multiple ancestry reference panels to further boost accuracy^17,18^. Singular value decomposition based approaches have also been proposed in other contexts^10^. In general, the accuracy of conditional association and imputation using summary statistics is reduced at low-frequency variants and when the LD structure between typed and imputed SNPs is mis-specified (e.g., when the ancestry of the GWAS sample does not exactly match the reference panel). We note that concerns about false-positive associations in imputation using summary statistics can be avoided entirely via the release of in-sample *summary LD information*, i.e. pairwise correlations between all typed SNPs.

### Gene-based association tests

#### Gene-based association using transcriptome reference data

GWAS risk variants are significantly enriched for genetic variants that impact gene expression (eQTLs)^21^. This motivates the paradigm of *transcriptome-wide association studies (TWAS)*, which evaluate the association between the expression of each gene and a complex trait of interest. Due to the limited availability of very large samples with measured gene expression and trait values, initial TWAS approaches integrated eQTL and GWAS to identify susceptibility genes either via matching the association signals^22-24^, via mediation analyses^25^, or via assessing whether the same causal variant impacts both gene expression and trait under a single causal variant model^26-28^.

More recent studies have leveraged predicted expression to improve the power of TWAS. Under this paradigm, transcriptome reference data is used to predict gene expression in the GWAS data set (using *cis* SNPs, e.g. within 1Mb of the transcription start site), followed by a test for association between predicted expression and trait. Although originally proposed using individual-level data^29^, TWAS using predicted expression can also be performed using only summary association statistics and summary LD information^30,31^. The key intuition is that the correlation between a weighted linear combination of SNPs (i.e. predicted gene expression) and trait is equivalent to a weighted linear combination of correlations between SNPs and trait (i.e. summary association statistics from GWAS) (see Figure 2). Since TWAS using predicted expression is conceptually similar to a test for non-zero genetic covariance between gene expression and trait^30^, it can also be performed via a two-sample *Mendelian randomization* from summary statistics^31^. TWAS using predicted expression can increase power over a standard GWAS when there exist multiple causal variants whose effect on trait is mediated through expression. TWAS also reduces the multiple hypothesis burden by testing tens of thousands of genes instead of millions of SNPs. TWAS using predicted expression typically uses individual-level transcriptome reference data to predict gene expression, but can also be performed using only summary association statistics between SNPs and gene expression, albeit with a reduction in power^30^. The potential power gains of TWAS are underscored by the recent identification of 71 new susceptibility genes across 28 complex traits, of which 17 have no GWAS association within 1 Mb^32^. However, TWAS is underpowered compared to standard GWAS when the true biological mechanism is independent of gene expression or when expression data in the most relevant tissue is not available.

**Figure 2.**
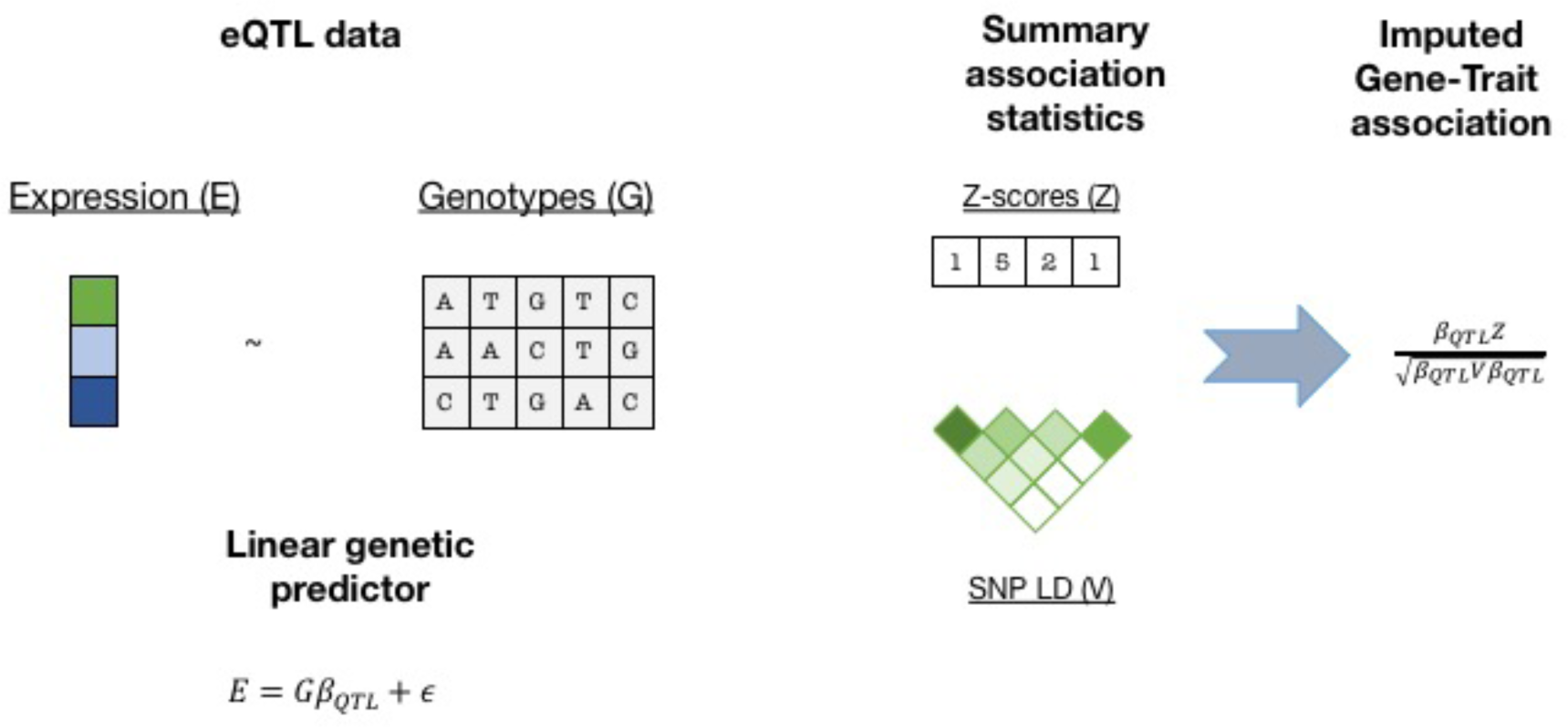
TWAS using predicted expression and summary data. TWAS using predicted expression and summary data follows two steps. First, transcriptome reference data is used to build a linear predictor for gene expression, typically using SNPs from the 1Mb local region around the gene with regularized effect sizes (e.g. using BSLMM^78^). Second, this predictor is applied to summary GWAS z-scores and gene-trait association z-scores are computed, testing the null model of no association between gene and trait.

#### Rare variant association tests

Although most GWAS of complex traits and diseases have focused on common variants that are typed on genotyping arrays or imputed from population reference panels, rare variant associations may also provide a rich source of biological insights, particularly for traits under strong negative selection^33,34^. Because association tests of individual rare variants are likely to be underpowered, rare variant association tests generally aggregate evidence for association across multiple rare variants at a locus. In exome sequencing studies (or exome array studies), rare variants are aggregated at the gene level, making the gene the unit of association. This can be done either using *burden tests*, which assume that all rare variants in a candidate gene have the same direction of effect, or using *overdispersion tests*, which assume that rare variants in a candidate gene can impact a complex trait in either direction; hybrid omnibus tests are also possible^35^. Recent studies have shown that both burden tests and overdispersion tests can be performed using only summary association statistics from each rare variant, together with summary LD information^36-38^ (see Box 2). Roughly, burden tests are computed as weighted sums of single-variant z-scores and overdispersion tests are computed as weighted sums of squared single-variant z-scores (analogous to previous work on common variant overdispersion tests using summary statistics^39^), with summary LD information used to specify appropriate null distributions in each case. However, a key limitation is that these studies require the use of in-sample summary LD information in preference to reference LD information to ensure appropriate null distributions and avoid false-positive associations. Thus, in contrast to summary statistic based methods for common variants (see above), both summary association statistics and in-sample summary LD information are required in order for these methods to be useful (see Discussion). An additional limitation is that, for case-control traits, asymptotic null distributions may not be valid when variant counts or case or control sample sizes are small, necessitating careful scrutiny of quantile-quantile plots.

### Fine-mapping

#### Fine-mapping using posterior probabilities of causality

Statistical fine-mapping aims to identify the causal variant(s) that are driving a GWAS association signal, enabling functional experiments to validate biological function. A straightforward approach to fine-mapping is to prioritize variants based on the strength of the marginal association statistics (i.e. ranking p-values)^40^. This is an effective strategy in the case of a single causal variant, but can be suboptimal when multiple causal variants are present, as the SNP with the top p-value at the locus may be tagging multiple causal variants. An alternative is to compute the posterior probabilities of causality for every SNP in the region, based on the likelihoods of the observed z-scores conditional on each possible set of causal variant(s)^41^. These posterior probabilities can be used to construct a credible set of SNPs, defined as the smallest set of SNPs that contains the true causal variant(s) with a given probability (typically 90% or 99%). Initial studies approximated the posterior probabilities of causality under a single causal variant assumption. Under this assumption, posterior probabilities of causality can be estimated from z-scores without the need for LD information^42^; this approach is both practical and computationally efficient. More recent studies have computed posterior probabilities of causality under a multiple causal variant assumption^43^. As in the case of imputation using summary statistics, the likelihoods of the observed z-scores can be computed based on the multi-variate normal (MVN) distribution with variance equal to the LD correlation matrix, with LD estimated from population reference panels using regularization techniques. Unlike imputation using summary statistics, which uses the null model of no association (i.e. a mean of 0 in the MVN), in fine-mapping the mean is a function of causal effect sizes, which can be heuristically approximated or integrated out using conjugate priors^43,44^. These methods often restrict computations to a maximum number of causal variants (e.g. 3 or 6); more recent studies have shown that further speed-ups can be achieved through matrix factorizations^45^ or stochastic search^46^. Methods that model multiple causal variants generally improve the accuracy (and calibration) of credible sets at loci with multiple causal variants^43-47^, with very limited decreases in accuracy at loci with only a single causal variant^43-49^. A less accurate alternative is to use conditional association analysis to detect multiple signals of associations^7,50,51^, followed by estimation of posterior probabilities of causality under a single causal variant assumption for each independent signal. In this case, special care is required in specifying the boundaries of each independent signal and the threshold for the conditional test.

#### Leveraging functional annotation data

Fine-mapping accuracy can be improved by integrating functional annotation data such as predicted regulatory elements from the ENCODE and ROADMAP Epigenomics projects^52,53^. This approach is motivated by early studies showing that disease-associated variants are systematically enriched in chromatin marks that delineate active regulatory regions in disease-relevant cell types^54,55^. Under this paradigm, a statistical model is developed to jointly estimate functional enrichment and update posterior probabilities of causality using functional annotations^44,49,56,57^. Some integrative methods assume that SNPs are unlinked^57^ or assume a single causal variant per locus^49,56^, but a recent study built upon the multiple causal variant model of ref. ^43^ to incorporate functional annotation data^44^. In an analysis of rheumatoid arthritis summary association data, integrative fine-mapping using this approach reduced the average size of 90% credible sets by 10%^58^. In addition to increasing fine-mapping accuracy, these studies have also provided insights into polygenic architectures (see below) by identifying tissue-specific functional annotations that are enriched for causal disease signals. This can also be achieved by conducting fine-mapping without integrating functional annotation data (typically under a single causal variant assumption) and then overlapping the resulting credible sets with functional annotation data to assess enrichment^59-61^. Future integrative methods could increase fine-mapping resolution by integrating probabilistic functional annotations (e.g., ChIP-seq peak intensity) or modeling the strength of association between SNPs and chromatin marks in population-based studies^62,63^.

#### Trans-ethnic fine-mapping

Fine-mapping accuracy can also be improved by leveraging differences in LD patterns across continental populations that have arisen due to differences in demographic events such as population bottlenecks (see Figure 3) ^64-67^. Intuitively, the set of tag SNPs linked to a causal variant will vary across populations, so that aggregating evidence of association across populations will dilute signals from tag SNPs and strengthen signals from causal variants. A standard approach to combining information across multiple studies is to compute posterior probabilities of causality from fixed-effects meta-analysis results^64,66,68,69^. Alternately, posterior probabilities can be computed from results of random-effects trans-ethnic meta-analysis methods^61,65^. These approaches assume a single causal variant and thus do not require LD information from the underlying populations. More recent studies have introduced hierarchical probabilistic models that allow for multiple causal variants while incorporating LD information from population reference panels^58^. These studies assume that causal variants are shared across populations but allow for heterogeneity in effect sizes across populations, and can also incorporate functional annotation data to further increase fine-mapping accuracy^58^. In an analysis of rheumatoid arthritis summary association data in Europeans and Asians (see above), trans-ethnic fine-mapping reduced the average size of 90% credible sets by 25%, and by 32% when also integrating functional annotation data^58^.

**Figure 3.**
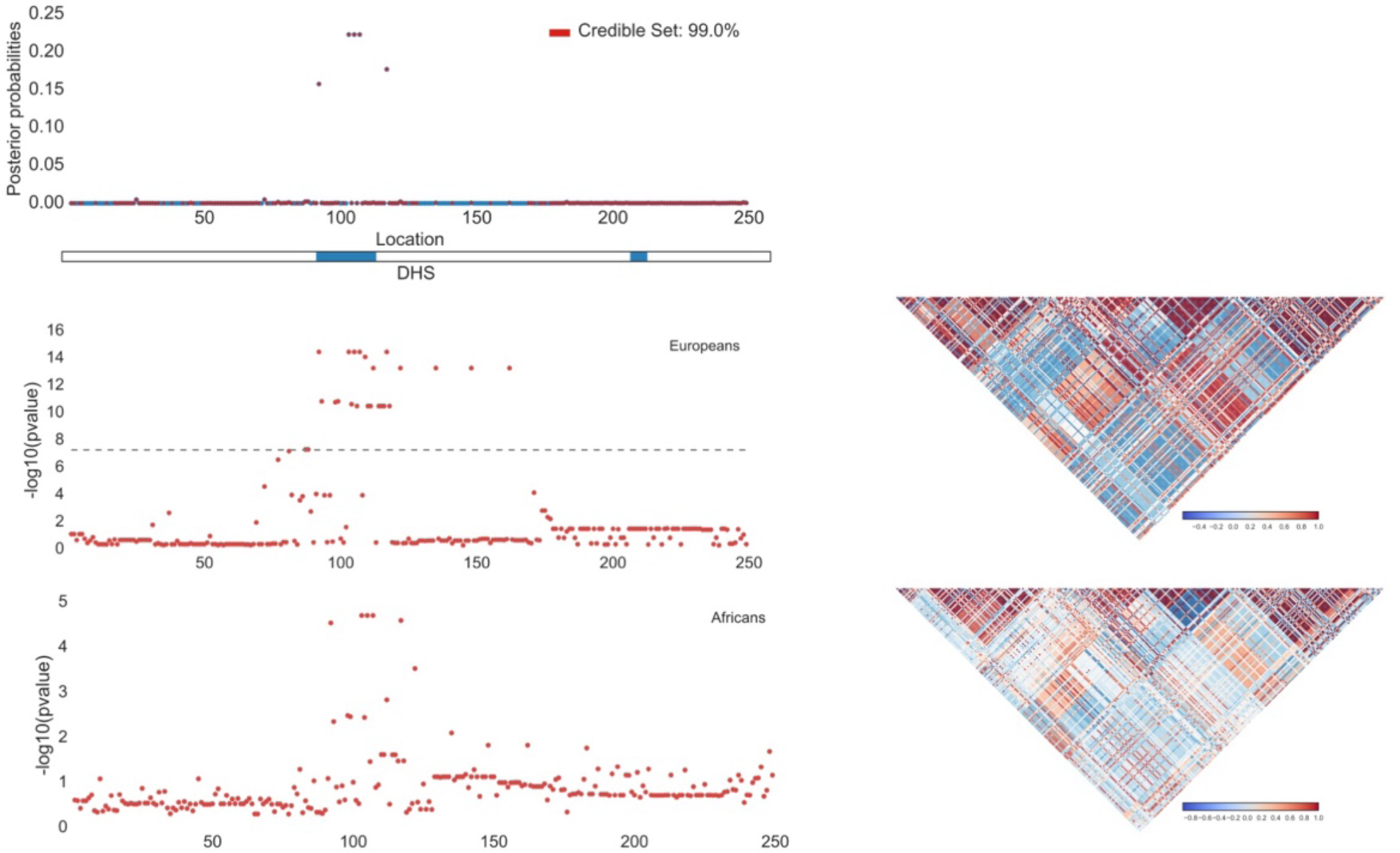
Leveraging functional annotation and trans-ethnic data to improve fine-mapping. A sample locus with simulated fine-mapping data in Europeans and Africans is displayed. The top panel shows the 99% credible set (denoted in red) produced by leveraging functional annotation data (DNase I Hypersensitivity Sites, DHS) in trans-ethnic fine-mapping. The middle and bottom panels show the –log 10 p-values (left) and LD (right) in Europeans and Africans.

### Polygenicity of complex traits

#### Polygenic risk prediction

Although the main focus of complex disease genetics is to gain insights about disease biology, genetics can also be leveraged to build predictions of disease risk, which may become clinically useful as sample sizes increase^70,71^. A landmark study of schizophrenia showed that *polygenic risk scores*, constructed by summing the predicted effects of all markers below a P-value threshold in the training sample, produced predictions of schizophrenia risk in validation samples that were significantly better than random, and far more accurate than those based on the single genome-wide significant locus identified in the study^72^. This provided an early demonstration of the advantages of incorporating markers that do not attain genome-wide significance into polygenic risk scores to improve prediction accuracy for polygenic traits. One complexity of polygenic risk scores is that of LD between markers, which has historically been addressed by LD-pruning—either without regard to P-values^72^, or via informed LD-pruning^73^ (clumping) that preferentially retains markers with more significant P-values. More recent work has shown that explicitly modeling LD using an LD reference panel and estimating posterior mean causal effect sizes can improve prediction accuracy from summary statistics^74^. An alternative to summary statistic based methods is to fit effect sizes of all markers simultaneously using Best Linear Unbiased Prediction (BLUP) methods and their extensions^75-77^, which require individual-level training data. Fitting all markers simultaneously is theoretically more appropriate and can produce more accurate predictions, although the relative advantage is small when overall prediction accuracies are modest (Box 3). In their simplest form, polygenic risk scores and BLUP methods assume infinitesimal (Gaussian) architectures in which all markers are causal, but these methods have been extended to increase prediction accuracy in the case of non-infinitesimal architectures; this has been accomplished for polygenic risk scores via restricting to markers below a P-value threshold^72^ or estimating posterior mean causal effect sizes under a point-normal prior^74^, and for BLUP methods by estimating (joint-fit) posterior mean causal effect sizes under a normal mixture prior^78,79^. Although polygenic risk scores must await even larger training sample sizes to attain clinical utility, appreciable prediction accuracies have been achieved for some traits, including a Nagelkerke *R*^2^ of 0.25 (AUC: 75%) for schizophrenia^74^. An important caveat is that it is critical when constructing and evaluating polygenic risk scores to avoid non-independence of training and validation samples (e.g. due to cryptic relatedness or shared population stratification), which could cause prediction accuracy to be overstated relative to what could be achieved in an independent validation sample^74,80^.

#### Inferring polygenic architectures

It is increasingly clear that most complex traits and diseases have highly polygenic architectures, with a large number of causal variants of small effect. In order to understand these polygenic architectures, it is of interest to infer parameters such as the heritability explained by SNPs and the number of causal variants. Both of these quantities have been estimated using accuracies of polygenic risk scores (see above), as a function of the P-value threshold used to constrain the set of markers employed^72,73^. Computing polygenic risk scores requires individual-level data in the validation cohort, implying that these methods are not strictly summary statistic based. Recent work has shown that the information in polygenic risk scores can be derived from summary-level data in the training and validation cohorts to estimate the heritability explained by SNPs and the number of causal variants^81^; a limitation of this approach is that SNPs are assumed to be uncorrelated, which can be approximately achieved by LD-pruning but precludes analyses of dense marker panels. The heritability explained by SNPs can alternatively be estimated from the slope of *LD score regression*^82^, in which *χ*^2^ statistics for each SNP are regressed against LD scores (sum of squared correlations with all SNPs), leveraging the fact that SNPs with higher LD scores are expected to contain more polygenic signal^83^. This approach explicitly allows for LD between SNPs and can distinguish between polygenicity and confounding, but makes strong assumptions about effect sizes of rare variants and thus currently only produces robust estimates for common variants. Another recent method models LD while treating SNP effects as fixed rather than random (similar to ref. ^81^), enabling estimation of heritability explained by common SNPs in local regions as well as genome-wide^10^. Overall, summary statistic based methods provide a useful alternative to methods for estimating heritability explained by SNPs from individual-level data using restricted maximum likelihood (REML) and its extensions^84,85^.

The increasing availability of functional annotation data (see above) can also be used to identify functional annotations that are enriched for polygenic signals of disease heritability. A recent study accomplished this using a Bayesian hierarchical model that splits the genome into blocks and incorporates both coarse-scale functional annotations at the level of blocks and fine-scale functional annotations at the level of SNPs^56^. This was the first study to quantify polygenic enrichments for cell-type-specific chromatin marks and DNase I hypersensitivity sites (DHS) across a broad set of complex traits and diseases. For example, polygenic signals for platelet volume and platelet count were enriched at DHS in CD34+ cells, which are on the cell lineage that lead to platelets, and polygenic signals for Crohn’s disease were depleted at repressed chromatin in LCL, an immune-related cell line. Functional enrichments can alternatively be estimated by stratified LD score regression^86^, which generalizes LD score regression^82^ to regress *χ*^2^ statistics for each SNP against LD scores with each functional category. Fine-mapping methods can also estimate functional enrichments, although these analyses are often restricted to disease-associated loci^44,49,58^. Notably, all of these summary statistic based methods have been applied to a large number of overlapping functional annotations, whereas methods that analyze individual-level genotypes have only been applied to a small number of non-overlapping functional annotations^87,88^. In addition, stratified LD score regression is not limited by the single causal variant per block assumption of the Bayesian hierarchical model, increasing power in settings of highly polygenic traits^86^. Application of the method identified significant cell-type-specific enrichments for many highly polygenic traits, including enrichments for histone marks in brain for smoking behavior and educational attainment—even though the summary statistics analyzed contained only one and three genome-wide significant loci, respectively. One limitation of the method is limited power for functional categories spanning a small percentage of the genome, motivating additional work in this area. As both summary statistic and functional annotation data sets grow larger and richer, identifying enriched functional annotations using summary statistic data will likely continue to be a fruitful endeavor.

### Cross-trait analyses

Many complex traits and diseases have a shared genetic etiology, either via shared genetic variant(s) with nonzero causal effect sizes (*pleiotropy*) or via a signed correlation between causal effect sizes (*genetic correlation*). Indeed, many instances of genetic variants with pleiotropic effects on multiple traits have been identified^89-94^. A recent study applied a Bayesian framework to summary association statistics from pairs of traits to estimate, at each locus in the genome, the probability that an associated variant has pleiotropic effects on both traits^95^. Pleiotropic SNPs can also be utilized as instrumental variables in Mendelian randomization analyses from summary statistics^96-98^, with one such analysis showing that increased body mass index causally increases triglyceride levels^95^.

An alternate approach to assessing the genetic overlap between two traits is to estimate the correlation between causal effect sizes across the two traits. Genome-wide genetic correlations can be estimated from individual-level data using bivariate REML^99^. A recent study estimated genome-wide genetic correlations from summary data using the information in polygenic risk scores, although this approach required LD-pruning the data which may lead to upwards bias^81^. Another recent study estimated genome-wide genetic correlations from summary data using cross-trait LD score regression^100^, which generalizes LD score regression to regress products of z-scores against LD scores for each SNP; this method produced estimates that were highly concordant with those from individual-level data^99^. Fitting the underlying MVN model using maximum likelihood instead of linear regression has produced promising results in applications to estimating cross-trait and cross-population genetic correlations, and may also prove useful in other settings^101^. Although genetic correlation analyses restricted to associated variants have also produced important findings^95^, the power of methods that leverage polygenic signals in genome-wide data is underscored by the discovery of significant genetic correlations involving traits with zero or few genome-wide significant loci, including a significant negative genetic correlation between smoking behavior and educational attainment^100^.

## Conclusion

Recently developed methods have made it possible to leverage summary association statistics to perform a wide range of analyses, many of which previously required individual-level data. As the availability of summary association statistics continues to grow (Table 1), summary statistics will continue to be broadly used in analyses involving single-variant association tests, gene-based association tests, fine-mapping, polygenic prediction and inferring polygenic architectures, and cross-trait analysis. The use of summary data will entail a slight loss of accuracy in some applications, such as imputation, where methods that analyze individual-level data can use haplotypes to model nonlinear structure, and polygenic prediction, where methods that analyze individual-level data can reduce noise by fitting all markers simultaneously; however, when summary statistics are available in larger sample size than individual-level data, the advantage of larger sample size will far outweigh those limitations. In addition, there are some settings where summary statistic based methods are the method of choice even when individual-level data is available, such as identifying functional annotations that are enriched for heritability, where methods that analyze individual-level data cannot currently handle a large number of overlapping annotations.

Despite considerable recent progress, there are some areas where further research on summary statistic based methods is needed. As population reference panels grow, more accurate modeling of rare and low-frequency variants will become possible, and it will be important to assess the limits of such efforts. It is also of interest to develop methods for inferring polygenic architectures from summary statistics that allow for different relationships between allele frequency and effect size. Identifying functional annotations that are enriched for heritability is an application that is particularly likely to produce important biological insights, and here there is a need for new methods that are well-powered for functional categories spanning a small percentage of the genome. As the number of functional annotations continues to increase, the integration of such data poses computational and statistical challenges in disentangling the correct functional annotations among many correlated ones.

We conclude by emphasizing the importance of making summary association statistics publicly available. A 2012 editorial in the journal *Nature Genetics* asked its authors to publish or database summary association statistics for all SNPs analyzed^102^, broadly impacting the set of publicly available summary statistics in the years that followed (Table 1). The public release of summary statistics is a useful compromise in situations where sample consent restrictions or privacy concerns preclude the release of individual-level data in a public repository. Although even the release of summary statistics can in principle lead to privacy concerns^103^, more recent work has shown that such privacy attacks have low power when the summary sample size exceeds the effective number of independent markers (currently estimated at 60,000 in typical GWAS data sets^104^), implying that privacy concerns should not preclude the public release of summary statistics from large studies^105-107^. Indeed, some recent studies have created web portals where summary data can be publicly accessed and visualized^60^. Finally, we note the potential benefits of publicly releasing summary statistics that include summary LD information (i.e. correlations) between each pair of proximal SNPs; however, the optimal approach to aggregating summary LD information across multiple cohorts in large-scale meta-analyses remains unclear, motivating future work in this area.

### Box 1: Conditional association and summary statistic imputation using LD reference data

Let *X* be an *N* x *M* matrix of genotypes, standardized to mean 0 and unit variance, and *Y* be an *N* x 1 vector of standardized trait values, where *M* is the number of SNPs at the locus and *N* is the number of samples. Under a standard linear model, *Y* = *Xβ* + *∊*. Let *V* be an *M* x *M* LD matrix of pairwise LD; *V* is equal to *X^T^X* if individual-level data is available, but can otherwise be estimated from a population reference sample (with or without regularization).

### Conditional association using LD reference data

We estimate the joint effects of all SNPs using least-squares as 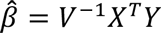 with 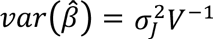, where 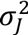 is the residual variance in the joint analysis. In a standard GWAS, however, each SNP is marginally tested one at a time, which can be expressed in matrix form as 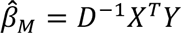 with 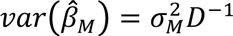, where *D* is the (nearly constant) diagonal matrix of *V* and 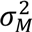 is the residual variance in the marginal analysis. It follows that:

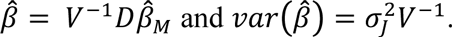

### Summary statistic imputation using LD reference data

Let 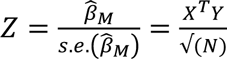 be a vector of z-scores (estimated effect sizes divided by their s.e.) obtained by marginally testing each SNP one at a time. Under the null hypothesis of no association, *Z* ∼ *N*(0,*V*). Let *Z_t_* and *Z_i_* partition the vector *Z* into *T* typed SNPs and *M* − *T* untyped SNPs, and let *V_t,t_* (covariances among typed SNPs), *V_i,i_* (covariances among untyped SNPs), and *V_t,i_* (covariances among typed and untyped SNPs) partition the matrix *V* accordingly. It follows that

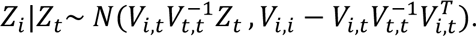

The mean and variance of the conditional distribution can be used to impute summary association statistics at untyped SNPs.

#### Box 2: Rare variant association tests using summary association statistics

Let *X* be an *N* x *M* matrix of genotypes, standardized to mean 0 and variance 1, and *Y* be an *N* x 1 matrix of standardized trait values, where *M* is the number of rare variants (e.g. in a given gene being tested for association) and *N* is the number of samples. An *M* x 1 vector of z-scores (estimated effect sizes divided by their s.e.) can be computed as 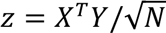, with multivariate normal null distribution *z* ∼ *N*(0,*V*), where *V* is an in-sample LD matrix.

### Burden tests

Burden tests assume that all rare variants in a candidate gene have the same direction of effect. Burden tests may either assume that standardized effect sizes are the same for each rare variant^108^ (i.e. per-allele effect sizes are proportional to 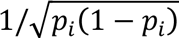, where *p_i_* is the allele frequency), or apply weights or thresholds based on allele frequency or functional information^109,110^. If *w* is an *M* x 1 vector of weights for each rare variant (including zero weights for rare variants excluded by a threshold), the test statistic for a weighted burden test is *T_burden_* = *w^T^Z* with null distribution *T_burden_* ∼ *N*(0, *w^T^Vw*). This test statistic can naturally be extended to meta-analysis of burden tests from multiple cohorts (via inverse-variance weighting), and can be extended to variable threshold tests and binary traits^36-38^.

### Overdispersion tests

Overdispersion tests assume that rare variants in a candidate gene can impact a complex trait in either direction, and can be computed as weighted sums of squared single-variant test statistics ^111,112^. If *W* = diag(*w*_1_,…, *w_M_*) is an *M* x *M* diagonal matrix of weights for each rare variant, the test statistic for a weighted overdispersion test is *T_overdispersion_* = *z^T^Wz* with null distribution 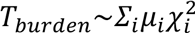, where weights *µ_i_* for each *χ*^2^ (1 d.o.f.) distribution 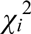 are given by eigenvalues of the matrix *V*^1/2^*WV*^1/2^. This test statistic can be extended to meta-analysis of overdispersion tests from multiple cohorts (via inverse-variance weighting), and can be extended to binary traits^36-38^.

#### Box 3: Polygenic risk prediction using summary vs. individual-level data

Suppose that polygenic risk prediction for a quantitative trait is conducted using a training cohort with *N* unrelated samples, using *M* unlinked markers with SNP-heritability^7^ equal to 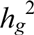. We initially consider two polygenic risk prediction methods that assume infinitesimal (Gaussian) architectures: polygenic risk scores computed using marginal effects at all markers with no P-value thresholding (PRS_all_), and fitting effect sizes of all markers simultaneously via Best Linear Unbiased Prediction (BLUP). We note that PRS_all_ requires only summary statistics from the training cohort, whereas BLUP requires individual-level data. The prediction *R*^2^ for each method are given by ^80,113^

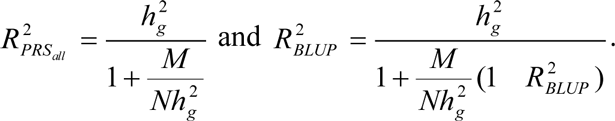

These equations can naturally be extended to linked markers (using the effective number of unlinked markers^104^) and case-control traits (using observed-scale SNP-heritability^114^). The relative advantage of BLUP over PRS_all_ is small when prediction *R*^2^ is small in absolute terms, but grows larger when prediction *R*^2^ is larger; this is illustrated in the figure below, which reports prediction *R*^2^ at various training sample sizes based on *M*=60,000 unlinked markers and a SNP-heritability of 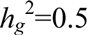. These results generalize to non-infinitesimal extensions of polygenic risk scores^72,74^ and BLUP^78,79^; in the latter case, the noise reduction from fitting all markers simultaneously remains equal to 1−*R*^2^, corresponding to an increase in training sample size of 1/(1−*R*^2^).

**Box 3 Figure:**

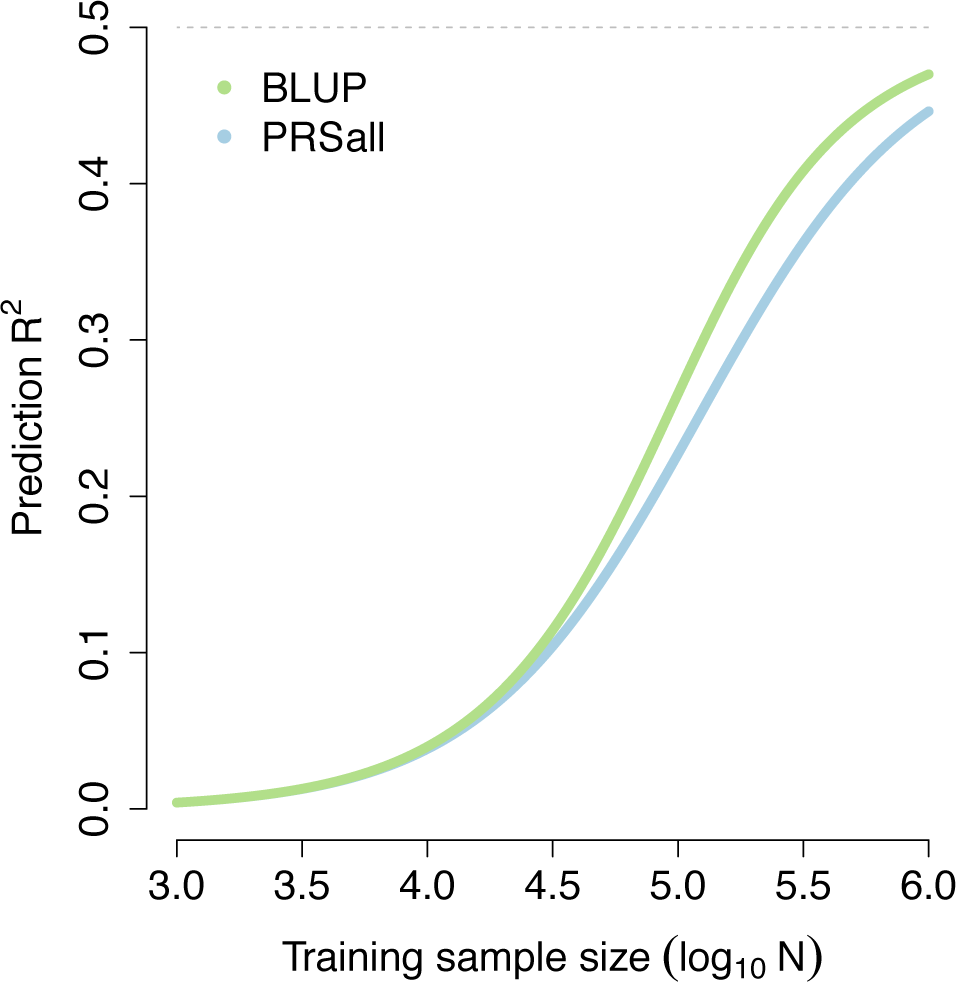

## Glossary

INDIVIDUAL-LEVEL DATA: Genome-wide SNP genotypes and trait values for each individual included in a GWAS.
SUMMARY ASSOCIATION STATISTICS: Estimated effect sizes and their standard errors for each SNP analyzed in a GWAS.
Z-SCORES: Association statistics that follow a standard normal distribution under the null; often computed as per-allele effect sizes divided by their standard error.
META-ANALYSIS: A method for combining data from different studies in which summary association statistics from each study are jointly analyzed.
MEGA-ANALYSIS: A method for combining data from different studies in which individual-level data from each study are merged and jointly analyzed.
SUMMARY LD INFORMATION: In-sample correlations between each pair of typed SNPs analyzed in a GWAS; can be restricted to proximal pairs of typed SNPs to limit the number of pairs of SNPs.
TRANSCRIPTOME-WIDE ASSOCIATION STUDY (TWAS): A study that evaluates the association between expression of each gene and a trait of interest; predicted expression may be used instead of measured expression to improve practicality.
MENDELIAN RANDOMIZATION: A method that uses significantly associated SNPs as instrumental variables to quantify causal relationships between two traits.
BURDEN TEST: A gene-based rare variant test in which all rare variants in a gene are assumed to have the same direction of effect.
OVERDISPERSION TEST: A gene-based rare variant test in which rare variants in a gene are assumed to impact trait in either direction.
POSTERIOR PROBABILITY OF CAUSALITY: The inferred probability that a SNP is causal, based on association data and optional prior information.
POLYGENIC RISK SCORE: A method of predicting trait by summing the predicted marginal effects of all markers below a P-value threshold in a training sample, multiplied by marker genotypes in a validation sample.
LD SCORE REGRESSION: A method of assessing trait polygenicity by regressing *χ*^2^ association statistics against LD scores for each SNP, computed as sums of squared correlations of each SNP with all SNPs including itself.
PLEIOTROPY: The existence of shared genetic variant(s) with nonzero causal effect sizes for two traits.
GENETIC CORRELATION: The signed correlation across SNPs between causal effect sizes for two traits.

## Acknowledgements

We are grateful to H. Finucane, S. Gazal, N. Mancuso and H. Shi for helpful discussions. We are grateful to G. Kichaev and R. Johnson for help with Figure 3. This work was funded by NIH grants R01 HG006399, R01 MH101244, R01 GM105857 and R01 MH107649.

## Key references

Ref. ^5^: This study introduced a powerful new random-effects meta-analysis method that employs a null model of no-heterogeneity.

Ref. ^7^: This study demonstrated that conditional association analysis can be performed using summary statistics.

Ref. ^12^: This was the first study showing that Gaussian imputation methods can be applied to summary-level genetic data.

Ref. ^26^: This study introduced a method for performing TWAS using summary statistics by assessing whether a single causal variant impacts both gene expression and trait.

Ref. ^30^: This study introduced a powerful method for performing TWAS using summary statistics by assessing the association between predicted gene expression (using all *cis* SNPs) and trait.

Ref. ^36^: This was the first of three studies demonstrating that rare variant burden and overdispersion tests can be performed using summary statistics.

Ref. ^42^: This study used posterior probabilities of causality to construct credible sets of causal disease-associated SNPs across multiple loci and diseases, under a single causal variant per locus assumption.

Ref. ^56^: This study used Bayesian hierarchical model to estimate posterior probabilities of causality and identify functional annotations enriched for disease heritability, under a single causal variant per locus assumption.

Ref. ^58^: This study showed that fine-mapping accuracy can be improved by leveraging functional annotation data and trans-ethnic samples and modelling multiple causal variants per locus.

Ref. ^72^: This study used polygenic risk scores to predict schizophrenia risk with appreciable accuracy, implicating a highly polygenic disease architecture.

Ref. ^95^: This study applied a Bayesian framework to identify pleiotropic effects across a broad set of complex traits and diseases.

Ref. ^100^: This study introduced a new method for estimating genome-wide genetic correlations from summary statistics.

